## Supplementary tables for "Transcriptome profiling of type VI secretion system core gene *tssM* mutant of *Xanthomonas perforans* highlights regulators controlling diverse functions ranging from virulence to metabolism"

A.

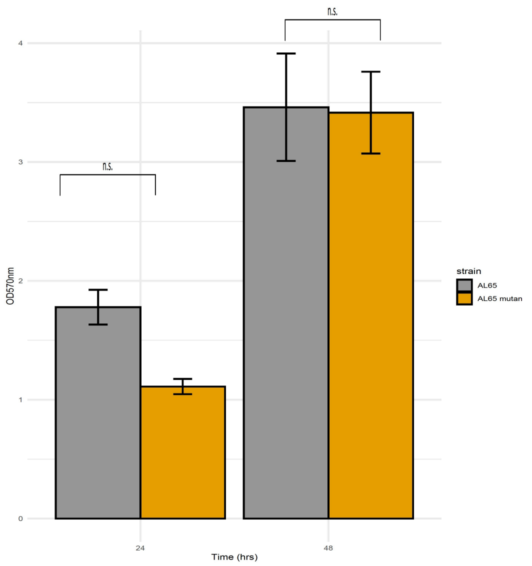


B.

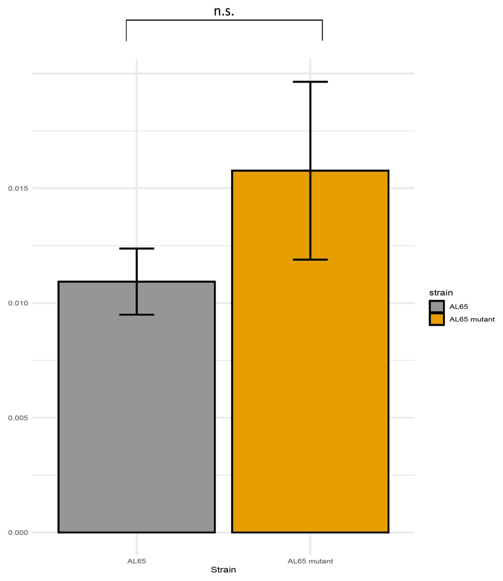


Fig S1. (A) Biofilm formation of AL65 and *tssM* mutant on abiotic surface quantified by measuring the absorbance (590 nm) of crystal violet. The strains were grown for 24 and 48 hours in XVM2 medium. The biofilm formed on the abiotic surface was stained with 0.1% crystal violet, excess stain was washed with dd$H_2O$ and air dried. The crystal violet stain was solubilized with 40% methanol and the absorbance was measured at 570 nm. The error bar indicates standard deviation. Eight biological replicates of each strain were performed. (B) EPS production by the wild type and mutant strain. The error bar indicates standard deviation. Three biological replicates of each strain were performed. n.s. denotes not significant.

Table S2 List of genes downregulated at 8h but upregulated at 16h in the *tssM* mutant strain.

| Locus tag | Gene | Product | Log2foldchange | |
|---|---|---|---|---|
| | | | 8h | 16h |
| E2P69_RS10240 | | PilN domain-containing protein | -0.90 | 1.27 |
| E2P69_RS00945 | | hypothetical protein | -0.66 | 1.24 |
| E2P69_RS02680 | *cheW* | chemotaxis protein | -0.84 | 1.00 |
| E2P69_RS17115 | | SDR family oxidoreductase | -0.73 | 0.98 |
| E2P69_RS13920 | | DUF4870 domain-containing protein | -0.63 | 0.97 |
| E2P69_RS08390 | | hypothetical protein | -1.26 | 0.90 |
| E2P69_RS01855 | | PilT/PilU family type 4a pilus ATPase | -0.69 | 0.89 |
| E2P69_RS17100 | | UDP-3-O-(3-hydroxymyristoyl)glucosamine N-acyltransferase | -0.62 | 0.88 |
| E2P69_RS02695 | | methyl-accepting chemotaxis protein | -0.86 | 0.86 |
| E2P69_RS17110 | | NeuD/PglB/VioB family sugar acetyltransferase | -0.60 | 0.86 |
| E2P69_RS02690 | | Hpt domain-containing protein | -1.04 | 0.86 |
| E2P69_RS17095 | | FkbM family methyltransferase | -0.70 | 0.82 |
| E2P69_RS17120 | | SDR family oxidoreductase | -0.75 | 0.81 |
| E2P69_RS02685 | | hypothetical protein | -0.95 | 0.80 |
| E2P69_RS10245 | *pilO* | type 4a pilus biogenesis protein | -0.83 | 0.80 |
| E2P69_RS10235 | *pilM* | pilus assembly protein | -0.80 | 0.79 |
| E2P69_RS17105 | | Rieske 2Fe-2S domain-containing protein | -0.68 | 0.79 |
| E2P69_RS17125 | | ketoacyl-ACP synthase III | -0.66 | 0.76 |
| E2P69_RS22030 | | GGDEF domain-containing protein | -1.81 | 0.74 |
| E2P69_RS10225 | | hypothetical protein | -0.65 | 068 |
| E2P69_RS10250 | *pilP* | pilus assembly protein | -1.02 | 0.62 |
| E2P69_RS08385 | | Cache 3/Cache 2 fusion domain-containing protein | -1.41 | 0.61 |
| E2P69_RS00330 | | TonB-dependent receptor | -0.61 | 0.60 |

Table S3 List of genes differentially expressed in the mutant strain associated with motility and chemotaxis

| Locus tag | Gene | Product | Log2foldchange | |
|---|---|---|---|---|
| | | | 8h | 16h |
| E2P69_RS17180 | *flgL* | Flagellin | -3.99 | -1.75 |
| E2P69_RS17255 | *flgM* | Flagellar biosynthesis anti-sigma factor | -3.96 | -2.47 |
| E2P69_RS17260 | *flgN* | Flagella protein | -3.78 | -2.16 |
| E2P69_RS17170 | *fliS* | Flagellar export chaperone | -3.67 | -1.48 |
| E2P69_RS22535 | *motD* | Flagellar motor protein | -3.60 | -1.89 |
| E2P69_RS22540 | - | Flagellar motor protein | -3.40 | -1.69 |
| E2P69_RS04575 | *motA* | Flagellar motor stator protein | -3.14 | -0.74 |
| E2P69_RS04570 | *motB* | Flagellar motor protein | -3.06 | -0.11* |
| E2P69_RS20890 | - | Flagellar brake protein | -2.84 | -0.89 |
| E2P69_RS22060 | *fliA* | RNA polymerase sigma factor | -2.61 | -0.79 |
| E2P69_RS17175 | *fliD* | Flagellar filament capping protein | -2.56 | -1.50 |
| E2P69_RS17185 | *flgL* | Flagellar hook-associated protein | -1.90 | -0.52* |
| E2P69_RS17190 | *flgK* | Flagellar hook-associated protein | -1.61 | -0.80 |
| E2P69_RS17195 | *flgJ* | Flagellar assembly peptidoglycan hydrolase | -1.45 | -0.88 |
| E2P69_RS17225 | *flgE* | Flagellar hook protein | -1.39 | -0.36* |
| E2P69_RS17220 | *flgF* | Flagellar basal-body rod protein | -1.26 | -0.26* |
| E2P69_RS17200 | *flgI* | Flagellar basal body P-ring protein | -1.24 | -0.45* |
| E2P69_RS17230 | *flgD* | Flagellar basal body rod modification protein | -1.13 | -1.08 |
| E2P69_RS22050 | *flhF* | flagellar biosynthesis protein | -1.11 | -1.79 |
| E2P69_RS22045 | *flhA* | Flagellar biosynthesis protein | -1.08 | -1.32 |
| E2P69_RS17205 | *flgH* | Flagellar basal body L-ring protein | -1.04 | -0.84 |
| E2P69_RS17035 | *fliN* | Flagellar motor switch protein | -1.00 | -0.35* |
| E2P69_RS17235 | *flgC* | Flagellar basal body rod protein | -1.00 | -1.82 |
| E2P69_RS17250 | *flgA* | Flagellar basal body P-ring formation protein | -0.99 | -1.93 |
| E2P69_RS17055 | *fliJ* | Flagellar export protein | -0.93 | -0.49* |
| E2P69_RS17030 | *fliO* | Flagellar biosynthetic protein | -0.86 | -2.04 |
| E2P69_RS17065 | *fliH* | Flagellar assembly protein | -0.86 | -0.34* |
| E2P69_RS17025 | *fliP* | Flagellar type III secretion system pore protein | -0.81 | -0.51 |
| E2P69_RS17050 | *fliK* | Flagellar hook-length control protein | -0.79 | -0.31 |
| E2P69_RS17070 | *fliG* | flagellar motor switch protein | -0.76 | -0.43* |
| E2P69_RS17075 | *fliF* | Flagellar M-ring protein | -0.73 | -1.31 |
| E2P69_RS17080 | *fliE* | flagellar hook-basal body complex protein | -0.58 | -1.90 |
| E2P69_RS17240 | *flgB* | flagellar basal body rod protein | -0.52* | -2.16 |
| E2P69_RS22040 | *flhB* | flagellar biosynthesis protein | -0.54* | -2.09 |
| E2P69_RS22010 | *fliQ* | flagellar biosynthetic protein | -0.24* | -2.08 |
| E2P69_RS17210 | *flgG* | flagellar basal-body rod protein | -0.54* | -1.90 |
| E2P69_RS17045 | *fliL* | flagellar basal body associated FliL family protein | -0.31* | -1.71 |
| E2P69_RS22015 | *fliR* | flagellar biosynthetic protein | -0.40* | -1.13 |

Table S4 List of genes differentially expressed in the mutant strain associated with pili biosynthesis.

| Locus tag | Gene | Product | Log2foldchange | |
|---|---|---|---|---|
| | | | 8h | 16h |
| E2P69_RS10895 | | molecular chaperone | -3.03 | -0.25* |
| E2P69_RS17160 | | PilZ domain-containing protein | -2.77 | -0.87 |
| E2P69_RS19765 | *pilA* | pilin PilA | -1.91 | 0.25* |
| E2P69_RS10240 | | PilN domain-containing protein | -0.90 | 1.27 |
| E2P69_RS10250 | *pilP* | pilus assembly protein PilP | -1.02 | 0.62 |
| E2P69_RS10255 | | type IV pilus secretin PilQ family protein | -0.83 | 0.40* |
| E2P69_RS10245 | *pilO* | type 4a pilus biogenesis protein PilO | -0.83 | 0.79 |
| E2P69_RS10235 | *pilM* | pilus assembly protein PilM | -0.80 | 0.79 |
| E2P69_RS02710 | *pilG* | twitching motility response regulator PilG | -0.75 | 0.04* |
| E2P69_RS01855 | | PilT/PilU family type 4a pilus ATPase | -0.68 | 0.89 |
| E2P69_RS01860 | *pilT* | type IV pilus twitching motility protein | -0.66 | 0.45* |
| E2P69_RS19730 | *pilB* | type IV-A pilus assembly ATPase PilB | -0.62 | -0.23* |
| E2P69_RS04995 | | PilT/PilU family type 4a pilus ATPase | -0.60 | 0.37* |
| E2P69_RS14450 | *pilE* | type IV pilin protein PilE | 1.45 | -0.08* |
| E2P69_RS14445 | | pilus assembly protein | 2.55 | 1.41 |
| E2P69_RS14440 | *pilX* | Tfp pilus assembly protein PilX | 2.77 | 1.64 |
| E2P69_RS14435 | | PilW family protein | 2.80 | 1.46 |
| E2P69_RS14430 | *pilV* | type IV pilus modification protein PilV | 2.96 | 1.33 |
| E2P69_RS10150 | | PilZ domain-containing protein | -0.22* | -0.72 |
| E2P69_RS21415 | *tadA* | Flp pilus assembly complex ATPase component TadA | 0.31* | 0.63 |
| E2P69_RS19760 | | pilin | -0.20* | 1.61 |

* Not significant

| | | | | |
|---|---|---|---|---|
| E2P69_RS17040 | *fliM* | flagellar motor switch protein | -0.49* | -0.96 |
| E2P69_RS18255 | | methyl-accepting chemotaxis protein | -4.85 | -1.74 |
| E2P69_RS18260 | *cheW* | purine-binding chemotaxis protein | -4.75 | -1.66 |
| E2P69_RS20895 | *cheW* | chemotaxis protein | -4.47 | -1.93 |
| E2P69_RS18250 | *cheA* | chemotaxis protein | -4.37 | -1.48 |
| E2P69_RS20845 | | MCP four helix bundle domain-containing protein | -4.32 | -1.14 |
| E2P69_RS20910 | *cheR* | chemotaxis protein | -4.23 | -1.70 |
| E2P69_RS09150 | *cheW* | purine-binding chemotaxis protein | -4.15 | -0.83 |
| E2P69_RS09155 | | MCP four helix bundle domain-containing protein | -4.08 | -1.19 |
| E2P69_RS20905 | | MCP four helix bundle domain-containing protein | -4.02 | -1.03 |
| E2P69_RS17295 | | chemotaxis protein | -4.00 | -1.84 |
| E2P69_RS22515 | *cheY* | response regulator | -4.00 | -1.29 |
| E2P69_RS22510 | *cheA* | chemotaxis protein | -3.95 | -0.97 |
| E2P69_RS12285 | | cache domain-containing protein | -3.53 | -0.02* |
| E2P69_RS18275 | | chemotaxis response regulator protein-glutamate methylesterase | -3.43 | -0.37* |
| E2P69_RS03320 | | methyl-accepting chemotaxis protein | -3.36 | -0.61 |
| E2P69_RS20920 | | chemotaxis response regulator protein-glutamate methylesterase | -3.33 | -0.04* |
| E2P69_RS22525 | *cheW* | chemotaxis protein | -3.17 | -0.93 |
| E2P69_RS20840 | | MCP four helix bundle domain-containing protein | -3.14 | -0.54* |
| E2P69_RS17245 | *cheV* | chemotaxis protein | -2.78 | -0.77 |
| E2P69_RS22065 | *cheY* | chemotaxis response regulator | -2.65 | -0.75 |
| E2P69_RS06725 | | PAS domain-containing methyl-accepting chemotaxis protein | -2.64 | -0.52* |
| E2P69_RS20915 | *cheD* | chemoreceptor glutamine deamidase | -2.63 | 0.04* |
| E2P69_RS22070 | *cheZ* | protein phosphatase | -2.61 | -0.69 |
| E2P69_RS22695 | *cheW* | chemotaxis protein | -2.22 | -0.65 |
| E2P69_RS20825 | | methyl-accepting chemotaxis protein | -2.19 | -0.11* |
| E2P69_RS20875 | | MCP four helix bundle domain-containing protein | -2.05 | -0.19* |
| E2P69_RS20870 | | MCP four helix bundle domain-containing protein | -1.88 | -0.41* |
| E2P69_RS09910 | | methyl-accepting chemotaxis protein | -1.68 | -0.37* |
| E2P69_RS02700 | *cheW* | chemotaxis protein | -1.03 | -0.52* |
| E2P69_RS02695 | | methyl-accepting chemotaxis protein | -0.86 | 0.86 |
| E2P69_RS02680 | *cheW* | chemotaxis protein | -0.83 | 1.00# |
| E2P69_RS11665 | *cheB* | chemotaxis protein | -0.74 | 0.11* |
| E2P69_RS11660 | *cheR* | chemotaxis protein | -0.73 | 0.07* |
| E2P69_RS20860 | | MCP four helix bundle domain-containing protein | -0.70 | -0.27* |

* Not significant

Table S5 List of genes differentially expressed in the mutant strain associated with c-di-GMP metabolism.

| Locus tag | Gene | Product | Log2foldchange | |
|---|---|---|---|---|
| | | | 8h | 16h |
| E2P69_RS18265 | | EAL domain-containing response regulator | -4.61 | -1.46 |
| E2P69_RS17280 | | EAL domain-containing protein | -4.05 | -1.49 |
| E2P69_RS06515 | | diguanylate cyclase | -3.51 | -0.97 |
| E2P69_RS12300 | | GGDEF domain-containing protein | -3.07 | -0.88 |
| E2P69_RS16355 | | GGDEF domain-containing protein | -3.03 | -0.62 |
| E2P69_RS13670 | | EAL domain-containing protein | -2.99 | 0.16[*] |
| E2P69_RS19000 | | EAL domain-containing protein | -2.63 | -0.37[*] |
| E2P69_RS21675 | | HD-GYP domain-containing protein | -2.27 | -0.53[*] |
| E2P69_RS22035 | | bifunctional diguanylate cyclase/phosphodiesterase | -2.15 | 0.06[*] |
| E2P69_RS12180 | | GGDEF domain-containing protein | -1.85 | -0.15[*] |
| E2P69_RS22030 | | GGDEF domain-containing protein | -1.80 | 0.74[#] |
| E2P69_RS10550 | | GGDEF domain-containing protein | -1.69 | 0.03[*] |
| E2P69_RS14210 | | EAL domain-containing protein | -1.18 | 0.01[*] |
| E2P69_RS17845 | | EAL domain-containing protein | -0.95 | 0.02[*] |
| E2P69_RS22020 | | diguanylate cyclase | -0.80 | -0.04[*] |
| E2P69_RS01315 | | GGDEF domain-containing protein | -0.67 | -0.62 |
| E2P69_RS10910 | | GGDEF domain-containing protein | -0.65 | -0.28[*] |

[*] Not significant

Table S6 List of differentially expressed genes in the mutant strain involved in type III secretion system.

| Locus tag | Gene | Product | Log2foldchange | |
|---|---|---|---|---|
| | | | 8h | 16h |
| E2P69_RS20480 | | HrpB1 family type III secretion system apparatus protein | -0.41* | -1.74 |
| E2P69_RS20475 | | type III secretion protein HrpB2 | -0.35* | -1.53 |
| E2P69_RS20500 | | type III secretion system cytoplasmic ring protein SctQ | -0.42* | -1.51 |
| E2P69_RS20495 | | type III secretion system protein SctP | -0.35* | -1.49 |
| E2P69_RS20535 | | CesT family type III secretion system chaperone | -0.45* | -1.49 |
| E2P69_RS20470 | | type III secretion inner membrane ring lipoprotein SctJ | -0.19* | -1.48 |
| E2P69_RS20465 | | type III secretion protein HrpB4 | -0.15* | -1.47 |
| E2P69_RS20505 | | type III secretion system export apparatus subunit SctR | -0.25* | -1.44 |
| E2P69_RS11735 | *hrpX* | helix-turn-helix transcriptional regulator | -0.14* | -1.41 |
| E2P69_RS11740 | *hrpG* | response regulator transcription factor | -0.15* | -1.39 |
| E2P69_RS20485 | | type III secretion system export apparatus subunit SctU | -0.38* | -1.38 |
| E2P69_RS20490 | | FHIPEP family type III secretion protein | -0.33* | -1.35 |
| E2P69_RS20555 | | HpaF protein | -0.16* | -1.33 |
| E2P69_RS20510 | | type III secretion system export apparatus subunit SctS | -0.30* | -1.30 |
| E2P69_RS20460 | | type III secretion system stator protein SctL | -0.07* | -1.29 |
| E2P69_RS20455 | | type III secretion system ATPase SctN | -0.17* | -1.27 |
| E2P69_RS20440 | | type III secretion system outer membrane ring subunit SctC | -0.24* | -1.25 |
| E2P69_RS20450 | | type III secretion protein HrpB7 | 0.01* | -1.21 |
| E2P69_RS20445 | | type III secretion system export apparatus subunit SctT | -0.07* | -1.67 |
| E2P69_RS04050 | | type III secretion system YopJ family effector AvrXv4 | -0.11* | -0.97 |
| E2P69_RS03165 | | type III PLP-dependent enzyme | 0.09* | -0.93 |
| E2P69_RS02615 | | type III secretion system effector protein XopK | 0.04* | -0.91 |

* Not significant

Table S7 List of type VI secretion system genes differentially expressed in the mutant strain.

| Locus tag | Gene | Product | Log2foldchange | | Cluster |
|---|---|---|---|---|---|
| | | | 8h | 16h | |
| E2P69_RS17690 | | type VI secretion system contractile sheath small subunit | -0.23* | 1.58 | i3* |
| E2P69_RS22610 | *hcp* | type VI secretion system tube protein | 0.33* | 1.46 | i3* |
| E2P69_RS12845 | | thiol:disulfide interchange protein DsbA/DsbL | 1.5 | 1.33 | |
| E2P69_RS08700 | | tetratricopeptide repeat protein | 0.53* | 1.10 | |
| E2P69_RS21880 | | type VI secretion system contractile sheath small subunit-cluster III | 0.00* | 1.04 | i3*** |
| E2P69_RS09395 | *tssK* | type VI secretion system baseplate subunit | 0.47* | 1.03 | i3* |
| E2P69_RS03325 | | thiol-disulfide oxidoreductase DCC family protein | 0.15* | 0.99 | |
| E2P69_RS21885 | *tssC* | type VI secretion system contractile sheath large subunit | 0.09* | 0.96 | i3*** |
| E2P69_RS09400 | *tssL* | type VI secretion system protein | 0.54* | 0.90 | i3* |
| E2P69_RS07570 | *tagH* | type VI secretion system-associated FHA domain protein | 0.25* | 0.88 | i3*** |
| E2P69_RS09410 | *tagF* | type VI secretion system-associated protein | 1.26 | 0.85 | i3* |
| E2P69_RS07575 | *tssK* | type VI secretion system baseplate subunit | 0.18* | 0.85 | i3*** |
| E2P69_RS21910 | *tssF* | type VI secretion system baseplate subunit | 0.23* | 0.80 | i3*** |
| E2P69_RS19505 | | tetratricopeptide repeat protein | 0.27* | 0.79 | |
| E2P69_RS05680 | | RHS repeat-associated core domain-containing protein | -0.17* | 0.78 | |
| E2P69_RS21920 | *clpV* | AAA family ATPase | -0.14* | 0.78 | i3*** |
| E2P69_RS09415 | | serine/threonine-protein phosphatase | 0.68 | 0.74 | |
| E2P69_RS21900 | | type VI secretion protein, ImpE/SciE family | 0.29* | 0.73 | i3*** |
| E2P69_RS09365 | | tetratricopeptide repeat protein | -0.10* | 0.73 | |
| E2P69_RS22605 | | ImpE protein | -0.07* | 0.72 | i3* |
| E2P69_RS21330 | | carbon storage regulator CsrA | 0.81 | 0.70 | |
| E2P69_RS09370 | *tssI* | type VI secretion system tip protein VgrG | 0.09* | 0.70 | i3* |
| E2P69_RS21905 | *tssE* | type VI secretion system baseplate subunit | 0.09* | 0.69 | i3*** |
| E2P69_RS14695 | | cardiolipin synthase B | 0.27* | 0.68 | |
| E2P69_RS17695 | *tssC* | type VI secretion system contractile sheath large subunit | 0.48* | 0.68 | i3* |
| E2P69_RS09360 | | PAAR domain-containing | 0.43* | 0.67 | i3* |

| | | | | | |
|---|---|---|---|---|---|
| | | protein | | | |
| E2P69_RS21890 | *hcp* | type VI secretion system tube protein | -0.06* | 0.66 | i3*** |
| E2P69_RS09440 | *tssA* | type VI secretion system protein | 0.27* | 0.66 | i3* |
| E2P69_RS17420 | | tetratricopeptide repeat protein | 0.48* | 0.65 | |
| E2P69_RS07585 | *tssM* | type VI secretion system membrane subunit | 0.31* | 0.61 | i3*** |
| E2P69_RS07625 | | two-component system VirA-like sensor kinase | -0.08* | 0.61 | |
| E2P69_RS09420 | | protein kinase | 0.20* | 0.59 | |
| E2P69_RS00975 | | phospholipase D family protein | 0.40* | 0.59 | |

* Not significant

**Table S8** List of genes associated with T2SS, T4SS and T5SS displayed differentially expression in the *tssM* mutant strain.

| Locus tag | Gene | Product | Log2foldchange | |
|---|---|---|---|---|
| | | | 8h | 16h |
| **T2SS** | | | | |
| E2P69_RS00220 | | glycoside hydrolase family 5 protein | -1.55 | -1.70 |
| E2P69_RS00225 | | glycoside hydrolase family 5 protein | -0.74 | -1.13 |
| E2P69_RS00235 | | glycoside hydrolase family 5 protein | 0.27* | -0.94 |
| E2P69_RS19010 | | cellulase family glycosylhydrolase | -1.05 | -0.03* |
| E2P69_RS02215 | | lipase | 0.08* | -0.82 |
| E2P69_RS15945 | | lipase | -0.15* | -0.78 |
| E2P69_RS05520 | | xylose isomerase xylA | 0.73 | 0.76 |
| E2P69_RS19555 | | xylanase | -0.32* | 0.86 |
| E2P69_RS05380 | | endo-1,4-beta-xylanase | 0.30* | -1.41 |
| E2P69_RS15315 | | alpha-amylase family protein | 0.40* | 1.02 |
| **T4SS** | | | | |
| E2P69_RS21420 | | TrbC/VirB2 family protein | 0.33* | 0.65 |
| E2P69_RS21425 | | VirB3 family type IV secretion system protein | 0.42* | 0.76 |
| E2P69_RS21460 | | TrbI/VirB10 family protein | 0.57* | 0.95 |
| E2P69_RS21450 | | type IV secretory pathway, TrbF protein | 0.58 | 0.84 |
| E2P69_RS21445 | | P-type conjugative transfer protein TrbL | 0.56* | 0.88 |
| E2P69_RS14500 | | TrbI/VirB10 family protein | -0.96 | -0.34* |
| E2P69_RS14505 | | P-type DNA transfer ATPase VirB11 | -0.94 | -0.04* |
| E2P69_RS14525 | | VirB4 family type IV secretion/conjugal transfer ATPase | -0.67 | -0.05* |
| E2P69_RS19420 | | P-type conjugative transfer protein TrbJ | -0.94 | -1.03 |
| E2P69_RS19425 | | TrbI/VirB10 family protein | -0.58 | -0.52* |
| **T5SS** | | | | |
| E2P69_RS19840 | | autotransporter domain-containing esterase | 0.95 | 0.47* |
| E2P69_RS22570 | | YadA-like family protein | 0.25* | 3.41 |
| E2P69_RS15045 | | YadA-like family protein | -0.03* | -0.94 |
| E2P69_RS17595 | | autotransporter-associated beta strand repeat-containing protein | -0.75 | -1.36 |
| E2P69_RS11440 | | DegQ family serine endoprotease | 0.17* | -1.32 |
| E2P69_RS16295 | | VirK family protein | -0.24* | -0.93 |
| E2P69_RS22135 | | preprotein translocase subunit SecE | 0.85 | 0.82 |
| E2P69_RS22465 | | preprotein translocase subunit SecY | 1.07 | 1.35 |
| E2P69_RS05570 | | Sec-independent protein translocase subunit TatA | -0.40* | -0.68 |
| E2P69_RS05575 | | twin-arginine translocase subunit TatB | -0.37* | -0.58 |

* Not significant

Table S9 List of differentially expressed genes coding for control of gene expression.

| Locus tag | Gene | Product | Log2foldchange | |
|---|---|---|---|---|
| | | | 8h | 16h |
| **Transcription, posttranscription and translational control of gene expression** | | | | |
| E2P69_RS17150 | *rpoN* | RNA polymerase factor sigma-54 | 0.17* | 0.88 |
| E2P69_RS21330 | *csrA* | carbon storage regulator CsrA | 0.81 | 0.70 |
| E2P69_RS10975 | | ribonuclease HII | 0.97 | 1.14 |
| E2P69_RS00065 | | ribonuclease P protein component | 0.64 | 1.04 |
| E2P69_RS10175 | | ribonuclease PH | -0.72 | -0.20* |
| E2P69_RS09245 | | rRNA maturation RNase YbeY | -0.65 | -0.56* |
| **Ribosomal Proteins** | | | | |
| E2P69_RS18785 | *rplL* | 50S ribosomal protein L9 | 0.85 | 0.59 |
| E2P69_RS10905 | | elongation factor Ts | 0.16* | 0.60 |
| E2P69_RS22455 | *rpmD* | 50S ribosomal protein L30 | 0.76 | 0.61 |
| E2P69_RS08540 | *rpmJ* | 50S ribosomal protein L36 | 0.89 | 0.61 |
| E2P69_RS07105 | | GTPase HflX | -0.14* | 0.62 |
| E2P69_RS00070 | *rpmH* | 50S ribosomal protein L34 | 0.90 | 0.62 |
| E2P69_RS22365 | *rplC* | 50S ribosomal protein L3 | 1.07 | 0.65 |
| E2P69_RS22405 | *rpmC* | 50S ribosomal protein L29 | 1.19 | 0.68 |
| E2P69_RS22370 | *rplD* | 50S ribosomal protein L4 | 1.04 | 0.74 |
| E2P69_RS22450 | *rpsE* | 30S ribosomal protein S5 | 0.85 | 0.75 |
| E2P69_RS17430 | | ribosome biogenesis GTPase Der | 0.44* | 0.79 |
| E2P69_RS11595 | *rplC* | 50S ribosomal protein L19 | 0.71 | 0.81 |
| E2P69_RS22425 | *rplE* | 50S ribosomal protein L5 | 0.70 | 0.82 |
| E2P69_RS22440 | *rplF* | 50S ribosomal protein L6 | 0.75 | 0.83 |
| E2P69_RS22375 | *rplW* | 50S ribosomal protein L23 | 0.93 | 0.84 |
| E2P69_RS22125 | *rplK* | 50S ribosomal protein L11 | 0.90 | 0.85 |
| E2P69_RS22395 | *rpsC* | 30S ribosomal protein S3 | 1.00 | 0.85 |
| E2P69_RS05925 | *rpmL* | 50S ribosomal protein L35 | 0.68 | 0.86 |
| E2P69_RS05930 | *rplT* | 50S ribosomal protein L20 | 0.79 | 0.86 |
| E2P69_RS22120 | | 50S ribosomal protein L1 | 0.83 | 0.90 |
| E2P69_RS22415 | | 50S ribosomal protein L14 | 0.93 | 0.92 |
| E2P69_RS22115 | | 50S ribosomal protein L10 | 0.95 | 0.92 |
| E2P69_RS21805 | | 50S ribosomal protein L33 | 1.05 | 0.93 |
| E2P69_RS03460 | *rpsU* | 30S ribosomal protein S21 | 1.10 | 0.95 |
| E2P69_RS22410 | *rpsQ* | 30S ribosomal protein S17 | 0.97 | 0.96 |
| E2P69_RS11610 | *rpsP* | 30S ribosomal protein S16 | 0.97 | 0.96 |
| E2P69_RS22390 | *rplV* | 50S ribosomal protein L22 | 0.91 | 0.97 |
| E2P69_RS11810 | *rpsT* | 30S ribosomal protein S20 | 1.08 | 0.99 |

| | | | | |
|---|---|---|---|---|
| E2P69_RS11605 | *rimM* | ribosome maturation factor | 0.66 | 1.00 |
| E2P69_RS14325 | *rimP* | ribosome maturation factor | 0.58 | 1.01 |
| E2P69_RS22400 | *rplP* | 50S ribosomal protein L16 | 0.89 | 1.01 |
| E2P69_RS22445 | *rplR* | 50S ribosomal protein L18 | 0.90 | 1.02 |
| E2P69_RS22430 | *rpsN* | 30S ribosomal protein S14 | 1.09 | 1.04 |
| E2P69_RS22385 | *rpsS* | 30S ribosomal protein S19 | 1.01 | 1.06 |
| E2P69_RS22490 | *rplQ* | 50S ribosomal protein L17 | 1.09 | 1.06 |
| E2P69_RS11820 | *rpmA* | 50S ribosomal protein L27 | 0.88 | 1.06 |
| E2P69_RS21800 | *rpmB* | 50S ribosomal protein L28 | 1.03 | 1.09 |
| E2P69_RS22380 | *rplB* | 50S ribosomal protein L2 | 1.16 | 1.11 |
| E2P69_RS22420 | *rplX* | 50S ribosomal protein L24 | 0.92 | 1.12 |
| E2P69_RS22460 | *rplO* | 50S ribosomal protein L15 | 0.88 | 1.13 |
| E2P69_RS22110 | *rplL* | 50S ribosomal protein L7/L12 | 0.99 | 1.13 |
| E2P69_RS22085 | | elongation factor G | 1.00 | 1.18 |
| E2P69_RS11825 | *rplU* | 50S ribosomal protein L21 | 0.91 | 1.19 |
| E2P69_RS14340 | | 30S ribosome-binding factor RbfA | 0.99 | 1.19 |
| E2P69_RS22345 | | elongation factor Tu | 1.15 | 1.48 |
| E2P69_RS11815 | | GTPase ObgE | 1.68 | 2.41 |
| E2P69_RS07135 | | ribosome assembly RNA-binding protein YhbY | -0.78 | -0.51* |
| E2P69_RS04970 | | 6-carboxytetrahydropterin synthase QueD | 1.17 | 0.53* |
| E2P69_RS11040 | | queuosine precursor transporter | 0.35* | -0.67 |
| E2P69_RS14330 | | transcription termination/ antitermination protein NusA | 0.46* | 1.06 |
| E2P69_RS22130 | | transcription termination/ antitermination protein NusG | 1.02 | 1.12 |

* Not significant

Table S10 Differentially expressed cell wall biogenesis and cell division genes in the mutant.

| Locus tag | Gene | Product | Log2foldchange | |
|---|---|---|---|---|
| | | | 8h | 16h |
| E2P69_RS06005 | | glycosyltransferase | 0.624 | 0.91 |
| E2P69_RS19735 | | glycosyltransferase family 2 protein | -0.36* | 0.92 |
| E2P69_RS06015 | | WecB/TagA/CpsF family glycosyltransferase | 0.24* | 0.95 |
| E2P69_RS05035 | | glycoside hydrolase family 99-like domain-containing protein | 0.37* | 1.00 |
| E2P69_RS19740 | | glycosyltransferase | 0.17* | 1.27 |
| E2P69_RS05995 | | glycosyltransferase | 0.57* | 0.81 |
| E2P69_RS05990 | | glycosyltransferase family 4 protein | 0.41* | 0.81 |
| E2P69_RS17085 | | glycosyltransferase | -0.39* | 0.84 |
| E2P69_RS07810 | | glycosyltransferase | 0.38* | 0.88 |
| E2P69_RS10970 | | lipid-A-disaccharide synthase | 0.84 | 1.10 |
| E2P69_RS03820 | | O-antigen ligase family protein | 0.77 | 0.62 |
| E2P69_RS05050 | | GtrA family protein | -0.37* | 0.76 |
| E2P69_RS22825 | | mannose-1-phosphate guanylyltransferase/mannose-6-phosphate isomerase | -0.07* | 0.79 |
| E2P69_RS08060 | | tetraacyldisaccharide 4'-kinase LpxK | 0.03* | 0.59 |
| E2P69_RS08055 | | 3-deoxy-manno-octulosonate cytidylyltransferase | 0.11* | 1.12 |
| E2P69_RS05980 | | acyltransferase family protein | 0.63 | 0.75 |
| E2P69_RS06020 | | gum cluster cupin domain | 0.64 | 0.33* |
| E2P69_RS05965 | | gumC family protein | 0.64 | 0.17* |
| E2P69_RS17100 | | UDP-3-O-(3-hydroxymyristoyl)glucosamine N-acyltransferase | -0.61 | 0.88 |
| E2P69_RS10955 | | UDP-3-O-(3-hydroxymyristoyl)glucosamine N-acyltransferase | 0.88 | 0.92 |
| E2P69_RS18425 | | UDP-N-acetylmuramoyl-L-alanine--D-glutamate ligase | 0.40* | 1.06 |
| E2P69_RS10965 | | acyl-ACP--UDP-N-acetylglucosamine O-acyltransferase | 1.08 | 1.12 |
| E2P69_RS17090 | | polysaccharide pyruvyl transferase family protein | -0.53* | 1.39 |
| E2P69_RS11805 | | murein biosynthesis integral membrane protein MurJ | 1.09 | 1.68 |
| E2P69_RS14115 | | transglycosylase SLT domain-containing protein | 0.62 | 0.72 |
| E2P69_RS14605 | | lytic murein transglycosylase MltB | 0.31* | 0.72 |

| | | | | |
|---|---|---|---|---|
| E2P69_RS14510 | | lytic transglycosylase domain-containing protein | -0.67 | -0.29* |
| E2P69_RS20410 | | lytic transglycosylase domain-containing protein MltE | -0.14* | -1.42 |
| E2P69_RS13385 | | FtsX-like permease family protein | 0.42* | 0.79 |
| E2P69_RS12520 | | rod shape-determining protein MreC | 0.99 | 1.01 |
| E2P69_RS12525 | | rod shape-determining protein MreD | 0.76 | 0.98 |
| E2P69_RS12535 | | rod shape-determining protein RodA | 0.01* | 0.60 |
| E2P69_RS17350 | | DNA translocase FtsK | 0.45* | 0.70 |
| E2P69_RS13080 | | cell division protein FtsL | 0.22* | -0.69 |
| E2P69_RS13125 | | cell division protein FtsQ/DivIB | 0.10* | -0.67 |
| E2P69_RS18205 | | cell envelope integrity protein CreD | 0.21* | -0.99 |

* Not significant

Table S11 List of genes associated with inorganic ion transport and metabolism.

| Locus tag | Gene | Product | Log2foldchange | |
|---|---|---|---|---|
| | | | 8h | 16h |
| E2P69_RS17150 | *rpoN* | RNA polymerase factor sigma-54 | 0.17* | 0.88 |
| E2P69_RS17540 | | NarK/NasA family nitrate transporter | -0.01* | 1.42 |
| E2P69_RS17550 | *nirD* | nitrite reductase small subunit | -0.15* | 1.60 |
| E2P69_RS17545 | *nirB* | NAD(P)/FAD-dependent oxidoreductase | -0.14* | 1.61 |
| E2P69_RS11645 | | response regulator NtrC family | -0.21* | 0.83 |
| E2P69_RS01055 | | nitrogen regulation protein NR(I) NtrC | -0.12* | 1.36 |
| E2P69_RS01040 | | P-II family nitrogen regulator | -0.08* | 1.07 |
| E2P69_RS15545 | | P-II family nitrogen regulator | 0.35* | 0.58 |
| E2P69_RS01045 | | ammonium transporter | 0.06* | 1.37 |
| E2P69_RS16795 | *phoR* | phosphate regulon sensor histidine kinase | 0.43* | 0.67 |
| E2P69_RS06675 | | PhoH family protein | 0.37* | 0.64 |
| E2P69_RS18535 | *pstS* | phosphate ABC transporter substrate-binding protein | 0.20* | 0.81 |
| E2P69_RS18520 | *pstB* | phosphate ABC transporter ATP-binding protein | 0.02* | 0.94 |
| E2P69_RS18515 | *phoU* | phosphate signaling complex protein | -0.05* | 1.03 |
| E2P69_RS18525 | *pstA* | phosphate ABC transporter permease | 0.12* | 1.09 |
| E2P69_RS18530 | *pstC* | phosphate ABC transporter permease subunit | 0.21* | 1.14 |
| E2P69_RS20335 | | histidine-type phosphatase | 0.58 | 1.28 |
| E2P69_RS18060 | | alkaline phosphatase family protein | -0.20* | 0.90 |
| E2P69_RS16410 | | glycerophosphodiester phosphodiesterase family protein | 0.43* | 0.99 |
| E2P69_RS15410 | *oprP* | porin | -1.00 | 0.21* |
| E2P69_RS02800 | *pqqB* | pyrroloquinoline quinone biosynthesis protein | -0.11* | -0.78 |
| E2P69_RS02810 | *pqqD* | pyrroloquinoline quinone biosynthesis peptide chaperone | -0.19* | -0.69 |
| E2P69_RS02805 | *pqqC* | pyrroloquinoline-quinone synthase | 0.03* | -0.67 |
| E2P69_RS02815 | *pqqE* | pyrroloquinoline quinone biosynthesis protein | 0.07* | -0.63 |
| E2P69_RS07775 | | YeeE/YedE family protein | 0.59 | 2.13 |
| E2P69_RS12595 | | sulfite exporter TauE/SafE family protein | 1.22 | 1.38 |

| | | | | |
|---|---|---|---|---|
| E2P69_RS07790 | | sulfite exporter TauE/SafE family protein | 0.67 | 0.71 |
| E2P69_RS13380 | | TauD/TfdA family dioxygenase | -0.16 | -1.08 |
| E2P69_RS07925 | | sulfurtransferase | -0.54 | -0.62 |
| E2P69_RS20745 | | cysteine synthase A | -0.14* | -0.74 |
| E2P69_RS20680 | *cysD* | sulfate adenylyltransferase subunit CysD | -0.09* | -0.61 |
| E2P69_RS21230 | | ferrous iron transport protein A | -0.11* | -0.70 |
| E2P69_RS10085 | | rubredoxin | -0.47* | -0.96 |
| E2P69_RS17575 | | ferritin-like domain-containing protein | -1.03 | 0.05* |
| E2P69_RS20330 | | siderophore-interacting protein | 0.62 | 0.67 |
| E2P69_RS00875 | | TonB-dependent siderophore receptor | 0.64 | 0.11* |

* Not significant

Table S12 List of dysregulated genes involved in amino acid and energy production.

| Locus tag | Gene | Product | Log2foldchange | |
|---|---|---|---|---|
| | | | 8h | 16h |
| E2P69_RS21115 | *hisB* | bifunctional histidinol-phosphatase/imidazoleglycerol-phosphate dehydratase | -0.13* | 1.17 |
| E2P69_RS21130 | *hisF* | imidazole glycerol phosphate synthase subunit | -0.21* | 1.20 |
| E2P69_RS21110 | *hisC* | histidinol-phosphate transaminase | -0.10* | 1.21 |
| E2P69_RS21120 | *hisH* | imidazole glycerol phosphate synthase subunit | 0.02* | 1.24 |
| E2P69_RS21135 | *hisIE* | bifunctional phosphoribosyl-AMP cyclohydrolase/phosphoribosyl-ATP diphosphatase | -0.57* | 1.27 |
| E2P69_RS21105 | *hisD* | histidinol dehydrogenase | -0.28* | 1.28 |
| E2P69_RS21125 | *hisA* | 1-(5-phosphoribosyl)-5-[(5-phosphoribosylamino) methylideneamino] imidazole-4-carboxamide isomerase | 0.19* | 1.59 |
| E2P69_RS21100 | *hisG* | ATP phosphoribosyltransferase | -0.09* | 1.62 |
| E2P69_RS21095 | | trp operon repressor | -0.28* | 1.75 |
| E2P69_RS21025 | *carB* | carbamoyl-phosphate synthase large subunit | 0.45* | 0.74 |
| E2P69_RS08780 | *argC* | N-acetyl-gamma-glutamyl-phosphate reductase | 0.46* | 1.17 |
| E2P69_RS08795 | *argE* | acetylornithine deacetylase | 0.24* | 1.24 |
| E2P69_RS08775 | *argH* | argininosuccinate lyase | 0.42* | 1.24 |
| E2P69_RS08790 | *argB* | acetylglutamate kinase | 0.23* | 1.28 |
| E2P69_RS07515 | *aroA* | 3-phosphoshikimate 1-carboxyvinyltransferase | 0.64 | 0.97 |
| E2P69_RS19925 | | 3-deoxy-7-phosphoheptulonate synthase | 0.63 | 0.72 |
| E2P69_RS16500 | | aminodeoxychorismate synthase component I | 0.82 | 0.62 |
| E2P69_RS04815 | | chorismate mutase | 0.75 | -0.19* |
| E2P69_RS09875 | | 3-isopropylmalate dehydrogenase | -0.48* | -0.64 |
| E2P69_RS09870 | | 3-isopropylmalate dehydratase small subunit | -0.53* | -0.97 |
| E2P69_RS09865 | | 3-isopropylmalate dehydratase large subunit | -0.60 | -0.80 |

| | | | | |
|---|---|---|---|---|
| E2P69_RS08130 | | succinate dehydrogenase, cytochrome b556 subunit | -0.72 | -0.97 |
| E2P69_RS08125 | | succinate dehydrogenase, hydrophobic membrane anchor protein | -0.69 | -0.93 |
| E2P69_RS08120 | | succinate dehydrogenase flavoprotein subunit | -0.61 | -0.75 |
| E2P69_RS08110 | | succinate dehydrogenase iron-sulfur subunit | -0.71 | -0.65 |
| E2P69_RS08725 | | cytochrome ubiquinol oxidase subunit I | -0.79 | -0.86 |
| E2P69_RS08195 | | c-type cytochrome | -0.69 | -0.84 |
| E2P69_RS13540 | | protocatechuate 3,4-dioxygenase subunit alpha | -0.33* | -1.41 |
| E2P69_RS21585 | | protocatechuate 3,4-dioxygenase subunit beta | 0.18* | -0.79 |

* Not significant

Table S13 Stress response genes differentially expressed in the mutant strain.

| Locus tag | Gene | Product | Log2foldchange | |
|---|---|---|---|---|
| | | | 8h | 16h |
| E2P69_RS16345 | | 1,4-alpha-glucan branching enzyme | 0.14* | 0.72 |
| E2P69_RS16335 | | 4-alpha-glucanotransferase | -0.07* | 0.73 |
| E2P69_RS16330 | *treY* | malto-oligosyltrehalose synthase | -0.02* | 0.60 |
| E2P69_RS16340 | *treZ* | malto-oligosyltrehalose trehalohydrolase | -0.03* | 0.69 |
| E2P69_RS12250 | *treA* | alpha,alpha-trehalase | -0.78 | -0.77 |
| E2P69_RS08795 | | acetylornithine deacetylase | 0.24* | 1.24 |
| E2P69_RS12820 | | betaine-aldehyde dehydrogenase | 0.43* | 1.52 |
| E2P69_RS15510 | *speA* | arginine decarboxylase | -0.25* | 0.66 |
| E2P69_RS12825 | *betT* | choline BCCT transporter | 0.70 | 1.38 |
| E2P69_RS08555 | | agmatine deiminase family protein | -0.00* | 0.62 |
| E2P69_RS09280 | *potB* | ABC transporter permease subunit | 0.41* | 0.85 |
| E2P69_RS09275 | *potC* | ABC transporter permease subunit | 0.21* | 1.34 |
| E2P69_RS06565 | *potA* | polyamine ABC transporter ATP-binding protein | 0.10* | 0.89 |
| E2P69_RS15800 | | CsbD family protein | 1.15 | 0.09* |
| E2P69_RS14465 | *uvrB* | excinuclease ABC subunit | 0.66 | 0.73 |
| E2P69_RS11830 | *uvrA* | excinuclease ABC subunit | 0.39* | 0.58 |
| E2P69_RS10375 | | OmpW family protein | -0.48* | -0.83 |
| E2P69_RS01815 | | OsmC family protein | -0.09* | -0.60 |
| E2P69_RS15710 | *rpoE* | RNA polymerase sigma factor | 1.05 | 1.48 |
| E2P69_RS09345 | *rpoE* | RNA polymerase sigma factor | -0.14* | 0.83 |
| E2P69_RS11450 | *rpoE* | RNA polymerase sigma factor | 0.29* | -1.00 |

* Not significant
